# Natural Variation in Preparation for Nutrient Depletion Reveals a Cost-Benefit Tradeoff

**DOI:** 10.1101/011148

**Authors:** Jue Wang, Esha Atolia, Bo Hua, Yonatan Savir, Renan Escalante-Chong, Michael Springer

## Abstract

Maximizing growth and survival in the face of a complex, time-varying environment is a common problem for single-celled organisms in the wild. When offered two different sugars as carbon sources, microorganisms first consume the preferred sugar, then undergo a transient growth delay, the “diauxic lag”, while inducing genes to metabolize the less preferred sugar. This delay is commonly assumed to be an inevitable consequence of selection to maximize use of the preferred sugar. Contrary to this view, we found that many natural isolates of *Saccharomyces cerevisiae* display short or non-existent diauxic lags when grown in mixtures of glucose (preferred) and galactose. These strains induce galactose-utilization (GAL) genes hours before glucose exhaustion, thereby “preparing” for the transition from glucose to galactose metabolism. The extent of preparation varies across strains, and seems to be determined by the steady-state response of GAL genes to mixtures of glucose and galactose rather than by induction kinetics. Although early GAL induction gives strains a competitive advantage once glucose runs out, it comes at a cost while glucose is still present. Costs and benefits correlate with the degree of preparation: strains with higher expression of GAL genes prior to glucose exhaustion experience a larger upfront growth cost but also a shorter diauxic lag. Our results show that classical diauxic growth is only one extreme on a continuum of growth strategies constrained by a cost-benefit tradeoff. This type of continuum is likely to be common in nature, as similar tradeoffs can arise whenever cells evolve to use mixtures of nutrients.

## Introduction

Natural environments contain complex, time-varying mixtures of nutrients and stresses. Understanding how cells use external cues to maximize growth and survival is key to understanding the evolution and function of regulatory circuits. Gene regulation allows cells to express pathways for specific tasks only in conditions when they are needed, to maximize the benefit of these pathways while minimizing their metabolic cost [1–4]. Regulatory circuits have evolved elaborate behaviors such as bet-hedging, signal integration, and environmental anticipation in response to the complexity of natural environments [5].

A classic example of gene regulation occurs during microbial growth on mixtures of carbon sources. For example, when budding yeast or *E. coli* grow in the sugars glucose and galactose, they first consume glucose, while dedicated signaling mechanisms repress galactose-utilization (GAL) genes [6–11]. When glucose has been exhausted, cells temporarily stop growing, induce GAL genes, and start growing again. The transient pause in growth, called the diauxic lag, can last up to several hours.

The diauxic lag is commonly thought to be a consequence of selection to minimize expression of superfluous metabolic pathways when a nutrient that can be more efficiently utilized is available [12–14]. This is supported by the observation that cells growing in two sugars that support similar growth rates do not exhibit a diauxic lag [8]. However, recent studies have shown that even in the same nutrient mixture, the length of diauxic lag can vary among experimentally evolved bacterial strains [15,16] or yeast isolates [17]. In both cases, evolved strains lacking a diauxic shift possessed weaker catabolite repression of secondary carbon pathways than the ancestor. This leads to a fitness cost during growth in the preferred nutrient, but a fitness advantage when the environment shifts rapidly between preferred and alternative nutrients. These results raise the question of whether similar mechanisms and fitness tradeoffs underlie the diauxic lag variation seen in natural yeast isolates [17].

To address this question, we monitored culture density and gene expression in ecologically diverse *Saccharomyces cerevisiae* natural isolates growing in mixtures of glucose and galactose. As expected, we found a spectrum of diauxic lag phenotypes, from strains with non-existent lags to those with more classical lag times of many hours. Strikingly, the variation in lag time is not due to differences in how fast strains can execute induction of GAL genes, but rather the timing of when they begin to induce. Short-lag strains induce GAL genes up to 4 hours before glucose is exhausted, in effect “preparing” for the transition to galactose metabolism. The degree of preparation correlates with the strength of glucose repression; strains that induce GAL genes at higher glucose levels also induce them earlier during diauxic growth. These results suggest that natural variation in catabolite repression is not only a key determinant of microbial fitness during sudden nutrient shifts [17], but also gradually changing nutrient conditions. Finally, we show that the observed phenotypic variation follows a tradeoff: early GAL induction benefits cells by preparing them for glucose exhaustion, but the cost of expressing GAL genes reduces growth rate while glucose is still present. This tradeoff is likely a general constraint on microbial growth strategies in mixed-nutrient environments.

## Results

### Natural yeast strains vary in length of diauxic lag

We grew 43 *Saccharomyces cerevisiae* strains in a carbon-limited medium containing 0.25% glucose and 0.25% galactose, the preferred and alternative carbon source respectively (Figure 1A). The *S. cerevisiae* strains come from a range of geographical locations and environments [18,19] and are all prototrophic, allowing us to omit amino acids from the media and avoid potential complications from their role as alternative carbon sources [20]. Bulk growth of the cells was measured by recording the optical density of each culture every 10 minutes for 44 hours using an automated plate reader (Materials and methods).

**Figure 1.**
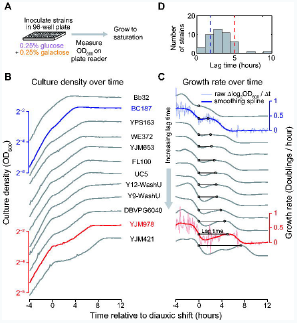
Natural yeast strains vary in length of diauxic lag. **(A)** Schematic of growth curve experiment in “diauxic growth conditions”, defined as batch culture in synthetic minimal media with 0.25% glucose and 0.25% galactose. **(B)** Growth curves (OD^600^ versus time relative to diauxic shift) plotted top-to-bottom in order of increasing diauxic lag. A single replicate growth curve is shown for each of 11 strains with similar growth rates in galactose-only media. (Growth curves for both replicates of all 43 strains assayed are shown in Figure S1). Strains BC187 and YJM978 are highlighted in red and blue, respectively. **(C)** Smoothed growth rate versus time relative to diauxic shift for the same strains as in (B). Example plots of raw OD differentials (light blue, light red) used to obtain the smoothed growth rate are shown for BC187 and YJM978. Diauxic lag time metric is denoted by horizontal black line with circles (see also Figure S2). (D) Histogram of diauxic lag time across all natural isolates assayed. Data used for histogram are the mean of 2 replicates (Materials and methods).

The growth curves generally display an initial phase of fast growth followed by a second phase of relatively slower growth, as expected in a 2-sugar mixture (Figure 1B–C, S1). However, the strains varied in the extent of growth lag, or a local minimum in growth rate, between the two growth phases (top versus bottom strains in Figure 1B–C). Some strains (e.g. YJM978) had a long diauxic lag during which growth rate almost reaches zero, whereas some strains (e.g. BC187) had a brief lag period during which even the minimum growth rate was relatively high. Strain SLYG78, a derivative of the commonly used laboratory strain S288C, exhibits a prominent lag phase (Figure S1), consistent with previous studies and the traditional understanding of *S. cerevisiae* as having a diauxic-growth phenotype [6,17]

To quantify the variation in diauxic lag, we defined a “diauxic lag time” metric as the time required to reach a strain’s smoothed maximal growth rate in galactose after having dropped below this growth rate during glucose depletion (horizontal black lines in Figure 1C, S2B, Materials and Methods). In growth curves that do not have a local growth-rate minimum, we defined the lag time as zero (Figure 1C, S2B). This lag time metric was robust to small differences in culture behavior (R^2^ = 0.96; Figure S2C) and to the method of calculation (Figure S2D, Materials and Methods).

We found that diauxic lag time varies continuously in our strains from 0 to 9 hours, with a mean of 3.2 hours and a standard deviation of 1.6 hours (Figure 1D). The continuous nature of the observed variation was robust to the choice of metric, as a related but distinct growth-curve feature, the minimum mid-diauxic growth rate, also varies continuously and correlates strongly with lag time (R^2^ = 0.71, Figure S2C). Lag time was not correlated with growth rate in pure glucose or galactose, and even among a subset of strains with similarly high growth rates in galactose-only media (subset shown in Figure 1B–C) we saw wide variability in the diauxic lag time (Figure S3). This suggests that the observed variation is due to differences in metabolic regulation rather than in maximal sugar utilization rates.

Several strains displayed no measurable diauxic lag and seem to transition instantly from glucose consumption to galactose consumption. This implies either that these strains can induce GAL genes quickly upon glucose exhaustion, induce GAL genes before glucose exhaustion, or both. To examine these possibilities, we characterized strains YJM978 and BC187, which represent long-lag and short-lag phenotypes, respectively (red and blue curves in Figure 1).

### Strain BC187 induces galactose-responsive genes before glucose is exhausted

We cultured BC187 and YJM978 in 0.25% glucose plus 0.25% galactose and monitored GAL pathway expression and glucose and galactose concentrations until saturation, when both sugars were depleted (Figure 2A, Materials and methods). We refer to this as a “diauxic growth experiment.” To enable single-cell measurement of GAL gene induction, we integrated a cassette containing yellow fluorescent protein driven by the GAL1 promoter (GAL1pr-YFP), which has been shown to be a faithful proxy for GAL pathway expression [21–23], at a neutral chromosomal locus (Figure 2A, top, Materials and Methods). We measured GAL1pr-YFP expression and extracellular sugar concentration by flow cytometry and enzymatic assay, respectively, over the entire diauxic growth cycle (Figure 2A, bottom). To quantify the timing of GAL gene induction, we defined *t*_low_ and *t*_high_, respectively, as the time when GAL1pr-YFP expression reaches 2-fold above basal levels and 1/4 of maximal levels, relative to the moment of glucose exhaustion (Figure 2B).

**Figure 2.**
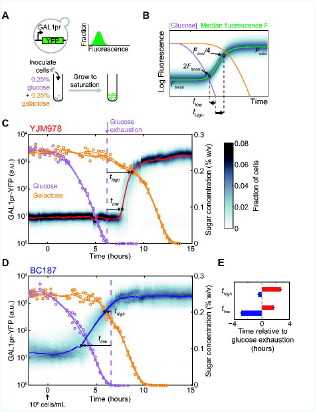
A short-lag strain induces galactose-utilization (GAL) genes hours before the diauxic shift. **(A)** *Top*: Schematic of GAL1pr-YFP transcriptional reporter and cartoon of fluorescence distribution as measured by flow cytometry. *Bottom*: Schematic of diauxic growth GAL induction experiment. **(B)** Definitions of induction metrics, *t*_low_ and *t*_high_, when reporter expression is at low but above-basal or near-maximal levels, respectively. Diauxic growth for strains **(C)** YJM978 and **(D)** BC187. GAL reporter expression distributions (gray shading), GAL reporter median (red line), glucose concentration (purple circles), and galactose concentration (orange circles). Time is defined relative to the moment when culture achieves a density of 10^6^ cells / mL (Figure S4). Purple and orange lines are smoothing spline fits to glucose and galactose measurements. Dotted purple line indicates time of glucose exhaustion, calculated using a local linear fit (Materials and methods). Data shown in (B) and (C) represent two replicate experiment. GAL reporter expression distribution is shown for only one of the two replicates. **(E)** Comparison of induction start time, *t*_low_, and near-maximal induction time, *t*_high_, for YJM978 (red bars) and BC187 (blue bars) cultures. Bars and error bars represent the mean and range, respectively, of two replicates.

Strain YJM978, which has a long diauxic lag, does not induce galactose-responsive genes until after glucose is exhausted, consistent with the classical understanding of diauxic growth (Figure 2C; *t*_low_ = 1.7 ± 0.1 hours, *t*_high_ = 2.7 ± 0.1 hours). In contrast, BC187, which has a short diauxic lag, begins GAL induction significantly before glucose exhaustion (Figure 2D; *t*_low_ = -3.0 ± 0.1 hours, p = 0.02 by t-test on n = 2 replicates). Even using the more conservative *t*_high_ metric, BC187 reaches near-maximal induction before glucose exhaustion (*t*_high_ = -0.5 ± 0.1 hours). Preinduction of GAL genes by BC187 leads to significant galactose consumption, even before glucose is fully exhausted (Figure S5). Both strains use glucose and galactose to completion and reach a similar yield (Figure S1), indicating that differences in induction time are not due to drastic differences in carbon utilization efficiency. Both strains have undetectable GAL1pr-YFP expression in glucose-only media (Figure S6), ruling out the possibility that galactose metabolism is constitutively active in BC187. In effect, BC187 “prepares” for the diauxic shift by inducing GAL genes before glucose exhaustion.Preparation is a continuous trait among natural yeast isolates

To determine if GAL induction prior to glucose exhaustion is a typical behavior of natural isolates, we integrated a GAL1pr-YFP reporter into 13 additional strains (for a total of 15 strains; see Table S1B) and monitored their expression during diauxic growth (same conditions as in Figure 2, Materials and Methods). Directly measuring sugar concentrations is laborious and less precise than measuring YFP fluorescence by flow cytometry, so we used YJM978 as a “reference” strain to signal the exhaustion of glucose, and co-cultured it with a “query” strain whose GAL induction kinetics we wanted to assay (Figure 3A). The reference strain was modified to express a fluorescent marker to distinguish it from the query strain (Figure S7A, Materials and Methods). To quantify differences in GAL induction time, we defined the “preparation time” as the difference in time between when the query and reference strains reach 1/16 of their maximal median GAL1pr-YFP expression (Figure 3B-C). Preparation time ranged from -3.8 to 0.04 hours relative to YJM978 with a mean of -1.3 hours, indicating that most strains induce GAL genes earlier than YJM978. The preparation time measured by this method is highly reproducible and robust to the query-to-reference mixing ratio (Figure S7B,C,E,F, Materials and Methods).If the degree of preparation determines the extent to which a strain has a diauxic shift, then strains that begin inducing GAL genes earlier should also have a shorter diauxic lag. We find a strong correlation (R^2^ = 0.83, p = 9.2×10^-7^) between preparation time and the diauxic lag time (Figure 3D). However, earlier-inducing strains appeared to take longer to reach full induction, or “execute” induction more slowly. We defined the “execution time” as the time required for median GAL1pr-YFP expression to increase from 1/64 to 1/4 of its maximal level (Figure 3B-C). The execution time anticorrelated with preparation time (Figure 3E inset) and lag time (Figure 3E), contradicting the naive expectation that a strain with a shorter diauxic lag will induce GAL genes more quickly. Taken together, our data show that the length of diauxic lag correlates to *when* strains begin to transition to galactose metabolism, not *how fast* they can execute the transition once they begin.

**Figure 3.**
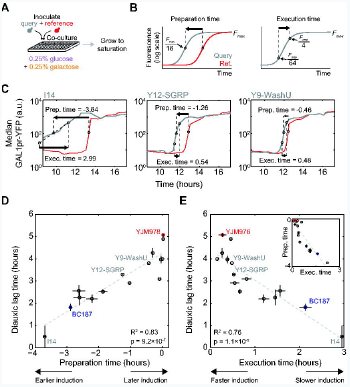
Diauxic lag time is correlated with the start time of GAL induction. **(A)** Schematic of co-culture GAL induction experiment. Each of 15 query strains (gray) are cocultured with reference strain YJM978 expressing constitutive mCherry marker (red), and sampled for flow cytometry every 15 minutes from mid-exponential phase until saturation. **(B)** Illustration of how preparation time and execution time metrics are defined. **(C)** Median GAL1pr-YFP expression versus time for query (gray) and reference (red) strain in 3 co-cultures selected to illustrate a range of preparation times. Strain I14 had above-basal reporter expression at the start of sampling, so its execution time was computed by linear extrapolation. **(D)** Scatterplot of diauxic lag time (from Figure 1) versus preparation time. **(E)** Scatterplot of diauxic lag time versus execution time. **(E, inset)** Scatterplot of preparation time versus execution time. Dotted gray lines in (D) and (E) indicate least-squares linear fits used to calculate coefficients of determination R^2^ and p-values. Data for diauxic lag time are the mean and range of two replicates, and for preparation time and execution time are the mean and s.e.m. of three replicates.

Related studies have observed population heterogeneity of growth rates and gene expression during sudden media shifts and diauxic growth [17,23,24]. In our experimental conditions, strains BC187 and YJM978 do not display bimodality in GAL1pr-YFP expression during diauxic growth (Figure 2C,D). A small number of strains do display bimodal expression at steady-state in glucose + galactose (Figure S6) and possibly also during diauxic growth (Figure S7D, G-I). However, the time window of any bimodality is short compared to induction time differences between strains (Figure S7G-I, Materials and methods). Therefore, although single-cell variation is likely an important dimension of regulatory behavior in some strains [17,23,24], our analysis of population-level dynamics already captures a major regulatory mode of diauxic growth.

### GAL induction kinetics after sudden media shift are poorly correlated with diauxic lag time

Our observations above rule out a model of the diauxic lag in which all strains begin inducing upon glucose exhaustion and vary only in how quickly they can reach maximal induction. However, it is possible that instead of inducing at glucose exhaustion, all strains induce when glucose is depleted below a certain threshold and vary in the delay before displaying observable GAL1pr-YFP expression. In this scenario, strains with a short delay between the start of induction and observable GAL1pr-YFP expression would appear to be preparing whereas strains with a long delay would appear to be inducing only after glucose exhaustion.

When cells are grown in glucose, the GAL pathway is repressed [7,25]. To ask whether differences in glucose de-repression kinetics could explain diauxic lag variation in our natural isolates, we grew strains in 2% glucose and transferred them into 2% galactose. We found significant variation in induction delay, defined as the time until median GAL1pr-YFP expression has increased 2-fold after transfer into galactose (Figure 4A). Some strains began to induce 5 hours after media switch, while one strain did not induce even after 18 hours. In contrast, the execution time of induction varied only between 0.6 to 1.6 hours (Table S2), suggesting that once glucose repression is relieved, GAL expression quickly induces from basal to maximal in all strains. Strikingly, induction delay after glucose-to-galactose shift was a poor predictor of both preparation time (Figure 4B; R^2^ = 0.16) and diauxic lag time (Figure 4C; R^2^ = 0.13). In particular, strains BC187 and I14 have short diauxic lags and early preparation times but very long induction delays after glucose-to-galactose media shift. Strain I14 had a similar behavior. When these two strains were omitted from the data, a weak correlation emerged (R^2^ = 0.56; p = 0.005), suggesting that glucose de-repression kinetics may play a role in the diauxic lag in our strains, but potentially convolved with a second response such as cell stress.

**Figure 4.**
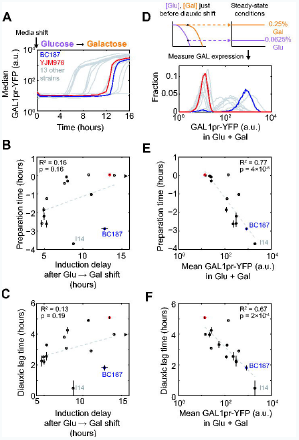
Diauxic lag time is correlated poorly with GAL induction kinetics but strongly with steady-state GAL expression in a glucose-galactose mixture. **(A)** Median GAL1pr-YFP expression versus time for BC187 (blue line), YJM978 (red line), and 13 other strains (gray lines) after transfer from 2% glucose into 2% galactose. **(B)** Scatterplot of preparation time (from Figure 3) versus induction delay after glucose-to-galactose shift, defined as the time until median GAL expression reaches 2-fold above basal expression. Black triangle indicates strain YJM981, which did not induce above background during the entire 18-hour experiment; this strain was omitted from the R^2^ calculation. (C) Scatterplot of diauxic lag time (from Figure 1) versus induction delay after glucose-to-galactose shift. **(D)** *Top:* Schematic of how sugar concentrations for steady-state measurements were chosen from the diauxic growth experiment. *Bottom:* Measured steady-state GAL1pr-YFP expression distributions for BC187, YJM978, and 13 other strains in 0.0625% glucose + 0.25% galactose. **(E)** Scatterplot of preparation time versus mean steady-state GAL1pr-YFP expression from (D). **(F)** Scatterplot of diauxic lag time versus mean steady-state GAL1pr-YFP expression from (D). Dotted gray lines in (B), (C), (E), and (F) indicate least-squares linear fits used to calculate coefficients of determination R^2^ and p-values.

### Differences in steady-state sugar sensing explains variation in preparation and diauxic lag time

Given that some strains can induce GAL genes in the presence of glucose, we hypothesized that differences in steady-state GAL expression in glucose + galactose may underlie differences in preparation. We measured GAL expression of our natural isolates in 0.0625% glucose + 0.25% galactose to simulate the conditions of a diauxic batch culture just before glucose exhaustion (Figure 4D). To ensure that we observed the steady-state response of the GAL reporter, we measured induction after cultures reached steady-state but before appreciable glucose had been depleted (Figure S8, Materials and Methods). We found that steady-state GAL expression in glucose + galactose varied from uninduced to almost maximal (Figure 4D, S6). On the other hand, all strains were uninduced in glucose-only media and maximally induced in galactose-only media (Figure S6), suggesting that strains vary not in overall glucose-repressibility or inducibility of GAL genes, but in how they integrate signals from both sugars in the mixed environment prior to diauxic shift.

We found that steady-state GAL expression in the glucose-galactose mixture correlated significantly with preparation time (Figure 4E; R^2^ = 0.77, p = 4×10^-5^) and diauxic lag time (Figure 4F; R^2^ = 0.67, p = 2×10^-4^). In other words, the strains that induce earlier during diauxic growth are those with higher steady-state GAL1pr-YFP expression in glucose + galactose. This suggests that short-lag strains do not suddenly switch GAL genes from “off” to “on” during diauxic growth, but instead express them at quasi-steady-state levels appropriate to the degree of glucose depletion. Consistent with this, the steady-state GAL1pr-YFP expression of these strains is proportional to their expression 3 hours before reference strain induction during diauxic growth (Fig S9A). Furthermore, BC187 grown in 3 sugar mixtures representing different moments during diauxic growth (0.25% galactose plus 0.25%, 0.125%, or 0.0625% glucose) displayed intermediate GAL1pr-YFP expression even after reaching steady-state, not basal or maximal expression as would be expected for a switch-like response (Figure S8, S9B). Taken together, our data strongly suggest that differences in preparation, and therefore diauxic lag time, are due to differences in the steady-state response of GAL genes to glucose-galactose mixtures.

### All strains prepare for glucose exhaustion during diauxic growth

Comparing the timing of GAL gene induction between diauxic growth and sudden media shift conditions offers a more sensitive measure of preparation for glucose exhaustion than the diauxic growth experiment alone. Even a long-lag strain like YJM978, which does not show observable GAL1pr-YFP expression until after glucose is exhausted (Figure 2 and 3), displays a much shorter induction delay during diauxic growth (*t*_low_ = 1.7 ± 0.1 hours; Figure 2C) than after media shift from glucose to galactose (induction delay = 12.2 hours, Figure 4A). To directly test if YJM978 could prepare for glucose exhaustion, we grew it in 0.125% glucose with or without 0.25% galactose and suddenly transferred the cells to galactose. We found that pre-growth in medium containing both galactose and glucose leads to an induction delay approximately one hour shorter than pre-growth in glucose alone, even though GAL1pr-YFP expression is indistinguishable from basal levels in both pre-growth media (Figure S10). As YJM978 has one of the longest diauxic lags in our set of strains, these data indicate that all strains prepare for glucose exhaustion to some degree.

### Preparation for glucose exhaustion has an immediate cost but delayed benefit

The fact that all of our strains prepared for glucose exhaustion by pre-inducing GAL genes suggests that preparation provides a fitness benefit. Consistent with this, strains with shorter diauxic lag times take less time after the diauxic shift to reach saturation (Figure 1B-C, S11A-B). But if preparation is always advantageous, then why don’t all strains display this phenotype? In the diauxic growth experiment of Figure 2, we noted that the YJM978 culture exhausts glucose 23 ± 4 minutes before BC187 does (Figure S11C), even though BC187 eventually exhausts both sugars first. Since BC187 and YJM978 grow at similar rates in glucose-only media (Figure S3), this suggests that BC187 is paying a cost for expressing GAL genes before glucose is exhausted.

To directly measure potential costs and benefits experienced by BC187 during diauxic growth, we performed a competitive fitness assay by co-culturing BC187 and YJM978 under diauxic growth conditions. In addition to GAL1pr-YFP reporter expression, we also monitored the relative abundance of the two strains by tagging them with constitutive fluorophores (Figure 5A, Materials and Methods). We plotted the log-ratio of BC187 to YJM978 cell counts versus time and found four different phases of relative fitness during a diauxic growth cycle (Figure 5A). Initially, when both sugar concentrations are high, both strains exhibit low GAL1pr-YFP expression (Figure 5B, Phase I) and grow at comparable rates (growth rate difference less than 0.062 doublings/hr at 95% confidence). When glucose is depleted below 0.1%, BC187 displays increased GAL1pr-YFP expression while YJM978 does not (Figure 5B, Phase II). During this phase, BC187 has a significant fitness disadvantage of -0.17 doublings/hr relative to YJM978 (Figure 5A, pink-shaded point in 5C; p = 0.0025 for non-zero slope by t-test). After glucose exhaustion, YJM978 begins to induce GAL1pr-YFP (Figure 5B, Phase III), and here BC187 has a significant fitness advantage of 0.38 doublings/hr relative to YJM978 (Figure 5A, light-blue-shaded point in 5C; p = 7.7×10^-5^ for non-zero slope by t-test). Once GAL1pr-YFP is fully induced in both strains the relative fitness again is comparable (Figure 5A, Phase IV; fitness difference less than 0.06 doublings/hour at 95% confidence).

**Figure 5.**
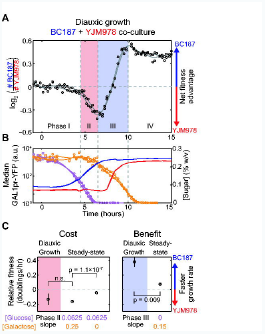
Preparation for glucose exhaustion has upfront cost and delayed benefit. **(A)** Log2-ratio of BC187 cell number versus YJM978 cell number versus time during diauxic growth in two replicate co-cultures. A positive value on the vertical axis at any given moment indicates that BC187 has divided more times than YJM978 since time = 0, and therefore has a net fitness advantage. Raw data (black circles) and smoothing splines (gray curves) are shown for two replicates. **(B)** Median GAL1pr-YFP expression of BC187 (blue lines) and YJM978 (red lines), glucose concentration (purple circles, lines), and galactose concentration (orange circles, lines) from (A). Definitions of “high”, “medium” and “low” sugar concentrations are indicated for reference. Vertical dotted gray lines in (A) and (B) demarcate 4 phases of relative fitness and GAL1pr-YFP expression during the experiment (see *Results*). **(C)** Comparison of growth rate differences during the diauxic growth versus steady-state sugar conditions. Data points with shaded backgrounds and labeled “diauxic growth” represent the slope of the data in (A) during Phase II (pink background) and Phase III (light blue background). Positive values indicate that BC187 grows faster than YJM978. Data are the mean and s.e.m. of n=6 (phase II) or n=14 (phase III) discrete derivatives in the shaded regions from (A). Points with white background and labeled “steady-state” are computed from the same data as in Figure S12C, and represent the mean and s.e.m. of 3-6 replicates. P-values are computed by 2-sample t-test.

This experiment shows that BC187 grows more slowly than YJM978 just before glucose exhaustion (Figure 5A, Phase II). To rule out that this is due to differences in utilization of low glucose concentrations unrelated to GAL regulation, we measured the absolute growth rates of the two strains in 0.0625% glucose with or without an additional 0.25% galactose,where sugar concentrations were held constant by frequent dilution (Figure S12, Materials and Methods). We found that BC187 grows at 0.62 doublings/hour in glucose alone, but significantly slower, at 0.51 doublings/hour, in glucose plus galactose (Figure S12C; p = 3.2×10^-4^ by t-test on n = 3-6 replicates per condition). YJM978 has the same growth rate of 0.67 doublings/hour in both conditions. Neither strain shows GAL1pr-YFP expression in glucose alone, but in glucose plus galactose, BC187 displays near-maximal induction while YJM978 remains at background (Figure S12D). These results correspond to a relative fitness of BC187 to YJM978 of -0.043 doublings/hour in glucose alone and -0.16 doublings/hour in glucose plus galactose. Only the latter is comparable to the fitness difference of -0.13 doublings/hour just prior to glucose exhaustion during diauxic growth (Figure 5C, left panel). Therefore, the fitness difference prior to glucose exhaustion is due to a *steady-state* cost of BC187’s early response to galactose.

In principle, the fitness difference after glucose exhaustion (Figure 5A, phase III) could be due to differences in galactose utilization rather than a benefit from pre-induction of GAL genes. To rule this out, we measured the steady-state relative fitness of the strains in 0.15% galactose (Figure S12C), corresponding to the carbon conditions just after glucose exhaustion when BC187 has its largest fitness advantage, 0.38 doublings/hour, over YJM978 (Figure 5A-B, Phase III),. In contrast, when galactose is held constant at 0.15%, BC187 has only a 0.076 doublings/hour advantage over YJM978 (Figure 5C, S12C). This steady-state relative fitness is significantly lower than the fitness difference during Phase III of diauxic growth (p = 0.009 by t-test; Figure 5C, right panel), showing that the majority of the fitness benefit after glucose exhaustion during diauxic growth is *kinetic*, not steady-state.

These results indicate that GAL pathway expression has a strong influence on growth rate in both constant and time-varying sugar environments. If this is a direct result of GAL gene activity, then cells from the same population with non-genetic variation in GAL expression should also exhibit different growth rates. To test this, we performed time-lapse microscopy to measure the growth rate and GAL expression of BC187 cells growing in 0.125% glucose + 0.25% galactose mixture, a partially inducing condition (Figure S13, Materials and Methods). To maximize the dynamic range of GAL expression of the observed cells, we performed three experiments, preinduced cells to low, medium, and high GAL1pr-YFP expression by culturing in 0.125% glucose, 0.125% glucose + 0.25% galactose, and 0.25% galactose respectively. We found that growth rate and GAL1pr-YFP expression displayed a significant negative correlation across cells of the same population, regardless of the pre-culture medium (Figure S13B). Furthermore, cell populations pre-induced to higher GAL1pr-YFP levels displayed lower growth rates than populations pre-induced to lower GAL levels (Figure S13C). Therefore, the fitness differences between bulk cultures of different strains may be due to effects of GAL expression at the single-cell level.

### Synthetic expression of GAL genes recapitulates costs and benefits

Given the long-established role of GAL genes in performing and regulating galactose metabolism [10], our findings strongly suggest that GAL expression causes the observed costs and benefits. Nevertheless, it is possible that unknown genes outside of the GAL pathway can also mediate cellular responses to the environments we studied. To show that expression of GAL pathway genes alone is sufficient to produce a fitness cost and a benefit, we introduced the chimeric transcription factor GEV into the S288C lab strain background (Figure 6A, “S288C-GEV”) [26,27]. The presence of β-estradiol, an otherwise inert compound in yeast, triggers the GEV protein to activate genes responsive to the GAL pathway activator GAL4p [28,29]. Therefore, S288C-GEV cells grown in glucose + β-estradiol will express all the inducible genes in the GAL pathway, as well as a GAL1pr-YFP reporter we integrated into this strain (Figure 6B, Materials and Methods). As expected, we find that S288C-GEV has a fitness cost relative to an unmodified S288C strain when grown in glucose + β-estradiol (Figure 6C, top panel, black line). This cost is absent in glucose-only media (Figure 6C, top panel, purple line), where S288C does not express GAL genes (Figure 6C, bottom panel).

**Figure 6.**
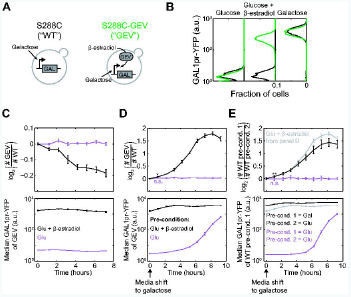
Synthetic induction of GAL genes is costly in glucose but beneficial during transition to galactose. **(A)** Strains S288C (“WT”) and S288C-GEV, a congenic strain expressing the GEV protein (“GEV”) were used. Both WT and GEV strains induce GAL genes (“GAL”) in response to galactose; strain GEV also induces GAL genes in response to β-estradiol. **(B)** GAL1pr-YFP expression histograms of strains WT (black) and GEV (green) at steady-state in 2% glucose, 2% glucose + 30nM β-estradiol, or 2% galactose. The same concentrations were used in the following experiments. **(C)** *Top*: log2-ratio of GEV to WT cell counts during steady-state coculture in glucose (purple) or glucose + β-estradiol (black). *Bottom*: median GAL1pr-YFP expression of strain WT during this experiment. **(D)** *Top*: log2-ratio of GEV to WT cell counts upon sudden shift to galactose, after pre-growth in glucose (purple) or glucose + β-estradiol (black). Asterisk ^“*”^: p = 0.008 for change in log2-strain ratio by 2-sample t-test. “n.s.”: not significant, or p > 0.05. *Bottom*: median GAL1pr-YFP expression of strain GEV during this experiment. **(E)** *Top*: log2-ratio of cell counts of two WT strains pre-grown in different conditions and shifted to galactose. The strains were either both pre-conditioned in glucose (purple) or the query strain (numerator of log-ratio) was pre-conditioned in galactose while the reference strain (denominator of log-ratio) was pre-conditioned in glucose (black). The black line from (D) is reproduced in gray in (E) to compare synthetic and “natural” GAL pre-induction. “**”: p = 0.01 for change in log2-strain ratio by 2-sample t-test. *Bottom*: median GAL1pr-YFP expression of the strain from pre-condition 1 during this experiment. The WT strain from pre-condition 1 contains a GAL1pr-YFP reporter, whereas the WT strain from pre-condition 2 expresses constitutive mCherry to distinguish the cells. Data in (C-E) are mean and s.e.m. of 3 replicates.

We find that S288C-GEV pre-induced in glucose + β-estradiol has an advantage over uninduced S288C when transferred suddenly to galactose medium (Figure 6D). We see a similar advantage when strain S288C is “naturally” pre-induced by growing in galactose, and then mixed with uninduced S288C and shifted together to galactose (Figure 6E). Therefore, induction of GAL genes recapitulates the benefits of galactose pre-growth in preparing cells for a transition to galactose. Surprisingly, the advantage of pre-induction (Figure 6D-E, slope of black line) is largest 3-6 hours after medium shift rather than immediately. However, this delay is seen for both synthetic and “natural” pre-induction, suggesting that it is due to stresses of the medium shift unrelated to sugar metabolism (Materials and Methods). In fact, even the immediate advantage of pre-induction is significant; by one hour after shift to galactose, the synthetically pre-induced strain has made 0.068 more doublings than the non-pre-induced strain (p = 0.008; “*” in Figure 6D). This is almost identical to the immediate advantage conferred by “natural” pre-induction (Figure 6E, gray and black lines). Therefore, expression of GAL genes alone is sufficient to cause a fitness cost in glucose-containing environments and a fitness benefit during transitions to galactose.

### Tradeoff between costs and benefits of preparation is a general constraint

Our data indicate that BC187 pre-induces GAL genes at a cost before the diauxic shift but reaps a benefit afterward, whereas YJM978 minimizes its preparation cost at the expense of experiencing a growth lag. To see if this tradeoff also constrains our other natural isolates, we assayed 15 strains to determine the cost they incur by responding to galactose while glucose is present. We defined the “galactose cost” of each strain as the relative difference in its steady-state growth rate in glucose + galactose versus glucose only, or specifically, as (*R*_glu+gal_ - *R*_glu_) / *R*_glu_, where *R*_glu+gal_ represents growth rate in 0.03125% glucose + 0.25% galactose and *R*_glu_ represents growth rate in 0.03125% glucose. Galactose cost ranged from 0 to -0.6, meaning that a strain may grow up to 60% slower simply because galactose is present in addition to glucose. The cost experienced by a given strain increased with its GAL1pr-YFP expression in glucose + galactose (Figure 7B), suggesting that the growth rate reduction is due to expression or activity of GAL genes. Additionally, when the cost measurement was repeated in 0.125% glucose + 0.25% galactose, a condition which elicits lower GAL1pr-YFP expression from most strains, the magnitude of galactose cost also decreased (Figure S14). These results confirm the presence of a tradeoff: no strain can partially induce GAL genes without also experiencing a decrease in growth rate.

**Figure 7.**
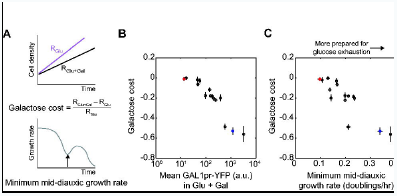
Tradeoff between costs and benefits of preparation underlies natural variation in GAL pathway expression. **(A)** Illustration of how (*Top*) galactose cost and (*Bottom*) the minimum mid-diauxic growth rate are defined (see also Figure S2, S14, and Materials and methods). Glucose and glucose + galactose conditions indicate 0.03125% glucose and 0.3125% glucose + 0.25% galactose media, respectively. **(B)** Scatterplot of galactose cost versus mean GAL1pr-YFP expression at steady-state in glucose + galactose. Data points are mean and s.e.m. of n=3 replicates. **(C)** Scatterplot of galactose cost versus minimum mid-diauxic growth rate. The latter is computed from the growth curves shown in Figure 1 and S1. Data points are the mean and s.e.m. of n=3 replicates (galactose cost) or mean and range of n=2 replicates (minimum rate).

To further illustrate this tradeoff, we used the minimum mid-diauxic growth rate (“minimum rate”) as a direct metric for the benefit of preparation (Figure 6A bottom). This metric is correlated with lag time and intuitively captures why preparation is beneficial: the more prepared a strain is, the higher its growth rate will be just after glucose exhaustion (Figure S2C). Furthermore, the minimum rate is not correlated with growth rate in glucose or galactose alone, and therefore is not convolved with steady-state metabolic differences (Figure S3). As expected, we found a negative correlation between preparation cost and minimum rate (Figure 7C). Our model strains for short-lag and long-lag phenotypes, BC187 and YJM978, appeared near the extremes of this tradeoff, with the phenotypes of most other strains in between.

## Discussion

### “Why no lag phase?” An old problem revisited again

A recent study by New et al. found that yeast strains evolved to respond quickly to sudden glucose-to-maltose (i.e. preferred-to-alternative sugar) transitions tended to also have shorter diauxic lags [17]. These evolved isolates acquired mutations that weakened carbon catabolite repression, so that maltose-utilization (MAL) genes are partially induced in otherwise repressing glucose levels. New et al. found that partial MAL expression is costly when glucose is available, but enables cells to resume growth more quickly when the environment changes suddenly from glucose to maltose.

Here we confirm the link between diauxic lag duration and glucose repression found by New et al., and observe an analogous expression cost in the GAL pathway, consistent with other reports [14,30]. Additionally, we extend the previous results in two ways. First, we show that variation in glucose repression leads to a spectrum of GAL pre-induction phenotypes during diauxic growth, and that this “preparation” is mediated by steady-state sugar-sensing rather than induction or de-repression kinetics. Secondly, we demonstrate that the same cost-benefit tradeoff that constrains lab evolution in sudden nutrient shifts also applies to natural isolates in gradually depleting nutrient mixtures. Overall, our results suggest that the mechanisms and selective forces that New et al. found in evolved strains are very likely also relevant in nature.

### Preparation arises from weak catabolite repression during gradual glucose depletion

Preparation for the diauxic shift can be attributed to two key features of the yeast GAL pathway. First, some strains express GAL genes at relatively high levels in glucose-galactose mixtures [31]. This partial induction has the effect of allowing cells to anticipate sudden nutrient shifts, which New et al. also hypothesized to underlie differences in diauxic lag duration [17]. However, it is not obvious *a priori* whether partial inducibility of GAL genes is physiologically relevant during diauxic growth, because glucose depletion must be slow relative to the timescale of response of GAL genes in order for significant pre-induction to occur. Our experiments demonstrate this second feature, and show that cells are indeed able to initiate (or continue) GAL induction during, and not only after, glucose depletion. For example, in our culture conditions, strain BC187 takes 4.1 hours and YJM978 3.3 hours to deplete glucose from 0.2% to 0%, while both strains can execute induction from 1/64 to 1/4 of maximal expression in less than 2 hours. Even long-lag strains, which do not display observable induction prior to the diauxic shift, still begin to induce sooner during diauxic growth than after a sudden nutrient shift, suggesting that all strains can prepare for glucose depletion. These findings contribute to growing evidence that batch culture is a continuous dynamical process and that this feature plays an important role in cellular regulation [32,33].

### Induction timing, not speed, underlies variation in diauxic lag

Previous studies have described differences in diauxic lag in terms of how quickly strains can transition from preferred to non-preferred nutrient metabolism [15–17]. We find that in a gradually depleting glucose-galactose mixture, “fast” or “slow” changes in growth rate are not due to “fast” or “slow” induction of GAL genes from basal to maximal, nor high basal induction, but rather “early” or “late” initiation of induction relative to glucose exhaustion. This clarifies a distinction between induction “speed” and “timing” that has not been addressed explicitly in previous work on diauxic growth.

New et al. observed a correlation between diauxic growth phenotype and growth delay after glucose-to-maltose shift [17], suggesting that common mechanisms underlie the behavior of cells in sudden and gradual nutrient shifts. We observe that diauxic lag duration is only weakly correlated to induction delay after a sudden glucose-to-galactose shift (Figure 4A-C), and instead that diauxic lag is more strongly correlated to preparation time and partial GAL expression (Figure 3, 4D-F). This discrepancy may be due to differences in our experimental systems, and suggest that our strains may experience stress after the glucose-to-galactose shift incurred by sudden loss of a metabolizable carbon source (Materials and Methods).

### Preparation as a widespread regulatory strategy

Other examples of preparation have recently been described in microbes encountering specific sequences of nutrients or stresses. For example, when *E. coli* encounter either heat shock or low oxygen, they induce both heat-responsive and low-oxygen-responsive genes, presumably an adaptive response when entering the warm, oxygen-deprived mammalian gut [34]. The co-regulation was decoupled by lab evolution under repeated heat shock in constant high oxygen, suggesting that the secondary response was neutral or costly when not needed. Anticipatory responses can also be asymmetric. When domesticated yeast encounter stresses typical of early stages of fermentation, they acquire resistance to later stresses; however, later stresses do not trigger resistance to early ones [35]. These results demonstrate that simple biochemical circuits can evolve the ability to anticipate environmental changes when the environmental cues occur in a predictable temporal sequence [36]. We now show that low or decreasing levels of a preferred nutrient can serve as a predictive cue for eventual depletion. Since this is inevitable when cells deplete a mixture of nutrients at unequal rates, and mixed-nutrient environments are ubiquitous in nature, environmental anticipation may be a more widespread regulatory strategy than previously recognized.

### Natural variation in diauxic lag may result from a tradeoff between costs and benefits

To be considered a meaningful example of preparation, a response must be beneficial in the future but neutral or costly in the present [35,36]. We showed that anticipatory GAL induction is costly--specifically, that many strains grow faster in glucose-only media than in media containing the same concentration of glucose plus an inducing concentration of galactose. The magnitude of cost is correlated to the degree of GAL expression across genetically diverse natural isolates, as well as across cells of the same strain with non-genetic expression variation. This cost can likely be attributed to the expression or activity of GAL pathway genes, because a strain that synthetically induces GAL genes in an otherwise non-inducing environment also exhibits a growth defect. These results rule out the possibility that strains induce GAL genes in glucose + galactose because it provides additional energy and thus a selective advantage.

The cost of GAL induction confirms part of the traditional rationale for the diauxic lag: strains that maintain stringent repression of alternative sugar pathways gain an advantage by maximizing their growth rate on glucose. On the other hand, we show that pre-induction also has a benefit that can sometimes more than compensate for its cost. Simply by being able to grow when glucose runs out, BC187 is able to double its population size over 3 hours while YJM978 undergoes a lag phase. This benefit is recapitulated when synthetically pre-induced cells are shifted from glucose to galactose media. The prevalence of short-lag phenotypes among natural strains shows that diauxic lag is by no means an inevitable phenotype in nature, and may be selectively advantageous only in certain conditions.

We find that strains seem to face a tradeoff between fast growth on glucose and readiness to grow on galactose when glucose runs out. In principle, these goals need not be in conflict, and given the countless ways that genetic variation can tune growth rates and gene expression, perhaps evolution can optimize multiple traits simultaneously. In fact, a naïve analysis reveals no tradeoff between our diauxic growth metrics and unnormalized growth rates in glucose or galactose (Figure S3), consistent with a similar observation by New et. al. [17]. Therefore, although the correlations that we observe across natural isolates suggest there can be a causal relationship between GAL gene regulation and fitness consequences during diauxic growth, definitive proof of this idea requires future work incorporating genetic and mechanistic analyses.

Given these caveats, it is nevertheless striking that we do observe a tradeoff between minimum diauxic growth rate and a galactose cost metric normalized for baseline growth rate differences in glucose. Like other examples of biological tradeoffs [2,3,37], our observation suggests the presence of underlying constraints despite substantial variation in other traits. In our strains, this constraint is likely the combination of an upper “speed limit” on how quickly GAL induction can be executed and an unavoidable cost of pre-induction.

### Bet hedging, mixed strategies, and optimal growth

In this study, we have focused on the timing of induction of entire cell populations during diauxic growth. Some of our natural isolates display bimodal GAL induction, similar to lab-evolved isolates, suggesting that the core phenomenon of preparation may be further modulated by heterogeneity across single cells. In fact, a different lab strain W303 has been found to implement both early and late induction strategies simultaneously in subpopulations of the same culture [23], reminiscent of microbial “bet-hedging” observed in other contexts [38–40]. This “mixed strategy” can be evolutionarily stable, as mutants with unimodal GAL induction are unable to invade the bimodal wildtype strain in glucose-galactose mixtures [41]. Similar population diversification during diauxic growth has been observed in bacteria [24,42,43]. Additionally, cellular decisions in nutrient mixtures can be influenced by epigenetic memory [22,44,45] and inter-species signalling [46,47]. An important goal of future investigation will be determining the relative importance of these different contributions to cellular decision-making in complex natural environments.

## Materials and Methods

### Strains and Media

Natural isolate yeast strains were obtained from multiple sources: 23 strains were part of the *Saccharomyces* Genome Resequencing Project (SGRP) and obtained from the National Collection of Yeast Cultures [18]; 18 strains were obtained from the Fay lab at Washington University [19]; strain Bb32 was obtained from the Broad Institute [48]; strain SLYG78 was constructed for this study. Some strains were obtained in duplicate, which we indicate by affixing “-SGRP” or “-WashU” to the strain name. One of these, Y12, displayed reproducibly different diauxic growth phenotypes depending on the source collection—this may be due to strain mislabeling (Table S1, personal communication, Justin Fay) [49]. All strains are homozygous diploid and prototrophic.

Growth curves were performed on 43 strains, and a subset of 15 natural isolates were chosen for subsequent analyses. A full strain list, as well as detailed genotypes of the 15-strain subset, can be found in Table S1. With the exception of SLYG78, the subset strains were transformed with vector SLVA63 or SLVD02 digested with NotI, which replaces the chromosomal HO locus with GAL1pr-YFP linked to the kanMX4 or hphNT1 selection marker respectively. Deletion of HO does not affect growth rate [50]. Strain SLYG78 was constructed by transforming S288C-lineage haploid strains FY4 and FY5 [51] with GAL1pr-YFP and TDH3pr-mTagBFP2 (vectors SLVD02 and SLVD13), respectively, and mating them to obtain a diploid. Strains BC187 and YJM978 were transformed a second time with SLVA19 or SLVA06, which replace the 2^nd^ HO locus with TDH3pr-mTagBFP or TDH3pr-mCherry linked to natMX4, respectively. These strains are designated BC187yb and YJM978ym in this section and in the supporting materials, but simply ‘BC187’ and ‘YJM978’ in the main text for clarity. Strain BC187ym was used for time-lapse microscopy experiments (Figure S13) instead of BC187yb (see “Single-cell measurements by time-lapse microscopy”); the two strains are identical other than the fluorescent protein they express. Strains for synthetic GAL induction via GEV are described below. All yeast transformations were done by the standard lithium acetate procedure [52].

All experiments were performed in synthetic minimal medium, which contains 1.7g/L Yeast Nitrogen Base (YNB) (BD Difco) and 5g/L ammonium sulfate (EMD), plus carbon sources. YNB contains no amino acids and extremely small amounts of other carbon-containing compounds, and therefore the added sugars comprise the sole carbon source. For diauxic growth experiments (Figures 1-3), the synthetic minimal media base was supplemented with 2.5g/L glucose (EMD) and 2.5g/L galactose (Sigma) to obtain 0.25% glucose plus 0.25% galactose w/v. We chose a 1:1 mixture of sugars to maximize the amount of growth curve data in both diauxic growth phases, and a total carbon concentration of 0.5% w/v because it is the highest that can be completely exhausted in synthetic minimal medium before non-carbon nutrients become yield-limiting. Unless noted otherwise, cultures were grown in a humidified incubator (Infors Multitron) at 30°C with rotary shaking at 230rpm (tubes and flasks) or 999rpm (deep 96-well plates).

### Growth curves and diauxic lag time metric

Growth curves (Figure 1) were obtained using an automated robotic workcell in a room maintained at 30°C and 75% humidity. Strains were cultured in 150uL of medium in optical-bottom 96-well plates (CellTreat). Plates were cycled between a shaker (Liconic) and a plate reader (Perkin Elmer Envision) using a robotic arm (Caliper Life Sciences Twister II), and absorbance at 600nm (OD_600_) was measured for each plate approximately every 10 minutes for up to 48 hours. In the humidity-controlled room, evaporation of medium was negligible within this time. Strains to be assayed were pinned from glycerol stock onto YPD agar and incubated for 16 hours, and then pinned into 600uL of liquid YPD and incubated another 16 hours. These cultures were diluted 1:200 into 600uL of synthetic minimal + 0.5% glucose and grown for 8 hours, and finally diluted 1:300 into synthetic minimal + 0.25% glucose + 0.25% galactose for growth curve measurements. The final inoculation was performed into 2 different plates; these replicate growth curves were nearly indistinguishable for all strains (Figure S1).

Analysis of growth curve data was performed in MATLAB using custom-written code. Raw OD_600_ readings were background-corrected by subtracting the median OD of 5-10 media-only wells on each plate. OD increased linearly with culture density in the density range of our cultures (Figure S2A). The OD of a typical saturated culture in our experiment was approximately 0.3, which corresponds to 5×10^7^ cells/mL. To analyze the diauxic lag, a smoothed growth rate was obtained by log2-transforming the data, computing discrete derivatives between consecutive data points as (<u>*OD*</u>_i_ – *OD*_i-1_) / (*t*_i_ – *t*_i-1_) and fitting the derivatives to a cubic spline using the MATLAB function `csaps` with a smoothing parameter of 0.75. This smoothed derivative represents the instantaneous growth rate in units of doublings/hour. The diauxic lag time metric was computed as the difference in time between the last local maximum in the smoothed growth rate and the previous point where the culture had the same growth rate; the earlier point was also used as the time of diauxic shift (Figure 1, S2B). The minimum mid-diauxic growth rate was computed as the minimum value of the smoothed growth rate between these two times (Figure S2B). In strains that did not have a local minimum in smoothed growth rate, we defined the diauxic lag as zero and the minimum mid-diauxic growth rate as the value of the smoothed growth rate at its inflection point between the two growth phases; this inflection point was also used as the time of diauxic shift (Figure S2B, strain Bb32). Similar results were obtained if the 2 metrics were calculated using a sliding-window average on the discrete derivatives instead of a smoothing spline (Figure S2D). We chose the smoothing-spline method because it facilitated calculation of a second derivative to allow identification of inflection points in the growth rate (Figure 2B, red lines).

To obtain growth rates in glucose or galactose (Figure S3), additional growth curves were performed as above, except the final culture medium contained 0.5% glucose alone or 0.5% galactose alone. Exponential growth rate was extracted from these data as the mean growth rate between when OD_600_ = 2^-6^ and OD_600_ = 2^-4^, or when culture density is approximately 1/16 and 1/4 of saturation.

### Flow cytometry and sugar assays on diauxic batch cultures

We assayed the gene expression and sugar consumption of BC187yb, YJM978yb, or a co-culture of the 2 during diauxic growth (Figure 2, 5) by inoculating them from single colonies into liquid YPD, incubating for 16 hours, mixing 1:1 by volume if co-culturing, and then diluting 1:100 – 1:500 into 2% raffinose and growing for 20 hours to ∼1.5×10^6^ cells/mL. The raffinose cultures were pelleted by centrifugation, washed once, and then resuspended in 0.25% glucose + 0.25% galactose medium in 2 replicate cultures of 50mL each. The cultures were incubated in flasks at 30°C with shaking, and a sample was removed every 15 minutes until saturation, about 18 hours. Some sample was placed on ice and diluted 1:2 – 1:100 in Tris-EDTA pH8.0 and read immediately on a Stratedigm S1000EX cytometer. The flow cytometer injected a defined volume, so we can estimate the absolute culture density (Figure S4A). The remaining sample was filter-sterilized using a Pall 0.2um filter plate and the flow-through stored at -20°C. Media flow-throughs were later thawed and assayed for glucose and galactose concentrations by mixing with a sugar-specific oxidase (Megazyme) and measuring the absorbance of the reaction at 340nm. A standard curve of known sugar concentrations was also assayed and used to infer concentration from absorbance. We expect YFP signal to change one hour slower than GAL1 protein levels, due to fluorophore maturation time [53]. This may be why galactose decreases slightly in the YJM978 culture before GAL1pr-YFP increases (Figure 2D). However, since all strains have the same reporter, this should not affect induction time differences between strains.

### Analysis of flow cytometry and sugar time course data

Flow cytometry data was analyzed using custom MATLAB code. In co-culture experiments, a 2D Gaussian mixture model (using the `gmdistribution` class) was fit to mCherry and side-scatter signal to segment the nonfluorescent and mCherry-expressing populations. When BC187yb was co-cultured with YJM978ym, segmentation was applied to both mCherry and BFP signal to exclude debris and doublets. We optimized flow cytometry conditions to minimize the occurrence of doublets (<1%), and therefore segmentation with 1 or 2 fluorescent markers gave equivalent results. GAL1pr-YFP expression histograms were computed on the log10-transformed YFP signal of each segmented subpopulation.

Results of diauxic growth experiments (Figure 2B-C, 5A-B) are plotted so that time zero corresponds to when culture density is 10^6^ cells/mL rather than inoculation time (Figure S4B-D). This allows the glucose consumption rate of each strain to be compared by looking at the glucose exhaustion time (Figure S11). To determine the glucose exhaustion time for each dataset in Figure 2, a line was fit to all glucose data points whose values lay between 0.01% and 0.05%, and the x-intercept of this line was taken as the time of glucose exhaustion. This method is more robust to measurement noise at low sugar concentrations than simply finding the time when concentration reaches some low threshold.

### Diauxic growth time-course measurements on multiple strains

To determine the timing of GAL pathway induction in multiple natural isolates (Figure 3), we co-cultured GAL1pr-YFP-labeled versions of each “query” strain with a “reference” strain, YJM978ym, which contains a constitutive fluorescent protein, TDH3pr-mCherry, as well as a GAL1pr-YFP reporter (Table S1; also see “Strains and media”). Query strains were grown in liquid YPD for 16 hours and then mixed with the reference strain YJM978ym at ratios of 1:4, 1:1, and 4:1 by volume. The mixed cultures were diluted 1:20 into YPD and grown for 4 hours, and then diluted 1:200 in 2% raffinose and grown for 12 hours. The raffinose cultures were then diluted 1:200 into 0.25% glucose + 0.25% galactose cultures split across 40 96-well plates. These were placed in a shaking incubator and allowed to grow for 8 hours before beginning sampling. Every 15 minutes a plate was removed from the incubator and its contents were mixed 1:1 with Tris-EDTA pH8.0 + 0.2% sodium azide to stop growth and protein synthesis, and incubated for 1 hour at room temperature to allow fluorophore maturation. The 40 plates were then measured on the flow cytometer with the aid of a robotic arm.

We confirmed that the constitutive fluorophore does not affect the time of induction by co-culturing two YJM978 strains, with and without the TDH3pr-mCherry (Figure S7B). We also compared the GAL induction start time (*t*_low_) of BC187 and YJM978 when they are cultured separately and when co-cultured, and saw no significant difference for either strain (Figure S7C). To check that growth rate differences between strains do not affect how quickly glucose is depleted, and therefore the timing of GAL induction, we performed each co-culture experiment at 3 different initial ratios of query to reference strain, and obtained almost identical results (Figure S7D-E). Therefore, this assay is robust to the presence and amount of reference strain, and we used the 3 inoculating ratios as replicates for data analysis.

To analyze population heterogeneity in GAL induction (Figure S7G-I), we computed the “ON fraction” as the fraction of cells with YFP signal greater than 1/32 of maximal median YFP. This threshold is just above the uninduced YFP level (Figure S7G). The ON fraction increases monotonically in most of our strains. Some strains have a small pre-induced population at the start of sampling (Figure S7H), consistent with the steady-state bimodality we have seen. Some strains do not seem to reach complete induction (ON fraction = 1), and in fact decrease in ON fraction due to an increasing YFP-off population toward the end of the timecourse (also see Figure S7D). This is unlikely for biological reasons (all glucose and most galactose has been depleted at that point) and may reflect the presence of contaminants in the fixative. Our metrics are computed on data before this potential contaminant reaches appreciable concentrations and do not affect the reported results. Likewise, before this point at least 90% of cells induce as one coherent population in all our strains (Figure 7H-I) rather than as two-subpopulations as seen by Venturelli et al. in strain W303 [23], which we did not assay here. The environmental and genetic determinants of induction time heterogeneity are potentially interesting to dissect in future experiments.

### Sudden medium shift experiments

The medium shift experiment in Figure 4A was performed by inoculating strains from colony into liquid YPD, incubating for 16 hours, and then diluting 1:500-8000 into 2% glucose so that cell density was approximately 1×10^6^ after 12 hours of further incubation. At this point, cultures were pelleted by centrifugation at 1000g for 2 minutes and washed once in 2% galactose. The cultures were pelleted again and resuspended in 2% galactose, and a sample of cells was removed from each culture and measured on the flow cytometer every 20 minutes for 18 hours. The same protocol was used when shifting strain YJM978ym from 0.125% glucose + 0.25% galactose to 0.125% glucose (Figure S10).

A similar experiment by New et al. using time-lapse microscopy after a glucose-to-maltose shift found that the average single-cell growth lag correlated with a metric similar to our diauxic lag time [17]. The apparent discrepancy between New et al. and our observations in Figure 4A-C is likely explained by differences in our metrics, the circuit studied (GAL versus MAL), and/or growth media. In particular, we used Yeast Nitrogen Base, which contains no carbon sources other than glucose or galactose, whereas New et al. used YP, which contains peptone and yeast extract. We speculate that auxiliary carbon sources may modulate the response of cells to sudden primary carbon shifts, a potentially interesting effect for future investigation.

### Calculation of induction metrics

For both the diauxic growth (Figure 2-3) and sudden medium shift experiments (Figure 4A), we analyzed GAL1pr-YFP expression kinetics by calculating the time that a certain threshold value of median YFP signal was reached, and using these induction times to define other metrics (e.g. preparation time). These induction time calculations were always done by linear interpolation between two data points that bracketed the threshold YFP value. The threshold values of YFP signal were chosen to reflect the meaning of a given metric—for example, we considered the “start” of induction to be when YFP signal reached 2-fold above the basal expression of that strain (usually the initial value in a timecourse), and the “end” of induction to be when YFP signal was 4-fold below maximal expression. If the same metric was used in different experimental designs (for example, execution time during diauxic growth or after medium shift), we occasionally chose different YFP thresholds to define the metric due to variation in the range of observed data. In general, however, our results were robust to the choice of threshold. For example, preparation time can be computed using a different definition of “mid-induction time” with almost identical results (Figure S7F). For a detailed description of each metric used in this study, and when they can be compared across experiments, see Table S2.

### Steady-state GAL expression and growth rate measurements

To measure the steady-state behavior of cells in defined glucose and galactose concentrations, we inoculated cells from colony into liquid YPD for 16 hours, diluted in 2% raffinose and grew for 20 hours, and then inoculated into glucose and/or galactose media and grew for at least 8 hours before sampling. To maintain steady-state conditions, we diluted the cultures 1:3 – 1:10 with fresh media every 2 hours so that the culture density stayed below 10^6^ cells/mL (Figure S12A, light-colored lines). Based on the observed glucose consumption rates, this ensures that less than 10% of the glucose in a 0.0625% glucose medium is depleted. As a further check, we continued the experiment up to 48 hours and found that GAL expression reached steady-state at 8 hours and stayed constant (Figure S8), indicating that our protocol was sufficient to prevent physiologically relevant changes in sugar concentrations.

To measure the steady-state relative and absolute growth rates of strains BC187yb and YJM978ym (Figure 5C,E), we co-cultured them in various glucose and/or galactose media and sampled and diluted the cultures every 2 hours for 12 hours. We determined the growth rate difference (a.k.a. selection rate) by fitting a line to the log2-ratio of cell counts for each strain over time (Figure 5C, S12B). We determined absolute growth rates from the same data by fitting a line to the log2-dilution-adjusted-cell-concentration (Figure S12A, C; see also Figure S4). We obtained precise dilution factors by weighing culture tubes when empty and during each dilution. These experiments were done with n=3-6 replicates. To compare steady-state growth rate differences to those from diauxic growth (Figure 5A), we computed discrete derivative of the log2-strain-ratio at all consecutive data points in Phase I or Phase II, and compared their distribution with our steady-state measurements by a 2-sample t-test (Figure 5C).

### Single-cell measurements by time-lapse microscopy

To prepare cells for time-lapse microscopy (Figure S13), we inoculated BC187ym from a colony into liquid YPD and grew for 16 hours, diluted in 2% raffinose and grew for 16 hours, and then diluted into 0.125% glucose, 0.125% glucose + 0.25% galactose, or 0.25% galactose for 8 hours to a density of 5×10^5^ cells/mL. Cells are then diluted 1:300 into 0.125% glucose + 0.25% galactose medium into wells (∼1000 cells / well) on a glass-bottom 96-well plate pre-coated with concanavalin A (Sigma) and left to settle for 1 hour. BC187ym contained a GAL1pr-YFP promoter and a TDH3pr-mCherry marker for image segmentation. Imaging was performed on a Nikon Eclipse Ti inverted microscope through a 20× objective lens. Exposures were taken every hour for 4 hours in bright field, YFP (ex. 500/24, em. 542/27), and mCherry (ex. 562/40, em. 641/75) channels, from 30 camera positions across 2 wells per pre-media condition, for a total of 90 camera positions. Image acquisition was controlled using custom MATLAB code using Micromanager/ImageJ.

Microscopy data were analyzed using custom MATLAB code. Microcolonies (clumps of 1-10 cells) were segmented in each mCherry image by applying a Gaussian blur to smooth cell boundaries, followed by a tophat filter to even out background, and thresholding to identify cells. Microcolonies were tracked across each time series by identifying overlapping areas. Colonies that split up, merged, entered, or exited the image during the acquisition time period were omitted from downstream analysis. Growth rate was computed as the change in log2 of a microccolony’s pixel area between first and last time points, divided by elapsed time (4 hours). YFP concentration was computed as the final average background-subtracted YFP signal per pixel area of a microcolony, where background YFP was taken as the median pixel intensity.

### Synthetic GAL induction using the GEV system

Synthetic induction experiments (Figure 6) were performed using 3 strains derived from FY5, a MATα S288C derivative (Table S1) [51]. Strain SLYA32 (“wt” reference strain in Figure 6C-E) was transformed with a constitutive TDH3pr-mCherry-natMX4 (vector SLVA06) to allow flow cytometry segmentation. Strain SLYA39 (“WT” in Figure 6B, query strain in 6E) was transformed with a GAL1pr-YFP-natMX4 reporter (vector SLVA64). Strain SLYH71 (“GEV” in Figure 6) was transformed with a tandem GAL1pr-YFP-ACT1pr-GEV-hphNT1 replacing the HO locus (vector SLVD04). The GEV sequence was subcloned from vector pAGL, a generous gift from the Botstein lab [26]. To perform competitive growth experiments (Figure 6C-D), query and reference strains were inoculated from single colonies into YPD, grown overnight, mixed 1:1 by volume, and then diluted 1:100 into YPD and grown 6 hours to OD∼0.3. Then the cultures were concentrated 5× by centrifugation and diluted in triplicate 1:300 (1:60 dilution of cells) into 2% glucose or 2% glucose + 30nM β-estradiol and grown 12 hours to pre-induce. If needed (Figure 6D), cells were shifted to 2% galactose by centrifugation at 3000g for 2min, washing in new medium, pelleting again, and resuspending. For the experiment in Figure 6E, the above protocol was used, except query and reference strains were kept in separate cultures until the time of medium shift, and then mixed and resuspended together into new medium. The cultures were sampled immediately after medium shift, and then every 30 minutes for 9 hours, to measure the strain ratio by flow cytometry. The query strain in Figure 6E, black line, is shifted from galactose medium back to the same medium, so the apparent delay in fitness advantage it exhibits may reflect a stress response to centrifugation and resuspension.

### Measuring galactose cost

To obtain the galactose cost (Figure 7, S14), we measured the growth rates of multiple strains in glucose and glucose + galactose. We co-cultured strains with the YJM978ym reference in 0.03125% glucose alone or 0.03125% glucose + 0.25% galactose (0.125% glucose in Figure S14), allowed them to grow for 8 hours, and then measured the cell count ratio at 2 timepoints 4 hours apart. To minimize glucose depletion, we inoculated cells so that their density at the end of the experiment did not exceed 3×10^6^ cells / mL. We computed the growth rate difference between query and reference strain as Δ*R* = [log_2_ (*N*_query,final_ / *N*_ref, final_) - log_2_ (*N*_query,initial_ / *N*_ref, initial_)] / 4 hours, where *N* refers to the number of cells of a particular strain at a particular timepoint. We computed the absolute growth rate of the reference strain in each well as *R*_ref_ = [log_2_ (*N*_query,final_ / *N*_ref, final_) - log_2_ (*N*_query,initial_ / *N*_ref, initial_)] / 4 hours, and then found the average and s.e.m. of reference strain growth rates across all wells of each condition as <*R*_ref,glu_> and <*R*_ref,glu+gal_> (see Table S2). We computed the absolute growth rates of query strains as *R*_query_ = <*R*_ref_ > + Δ*R* in each of the two conditions. Then we computed the galactose cost metric as (*R*_glu+gal_ - *R*_glu_) / *R*_glu_, where *R* denotes query strain growth rates in each condition. Error bars are the s.e.m. of galactose cost, computed from the s.e.m. of measured Δ*R* and <*R*_ref_ > values using standard uncertainty propagation formulas [54].

### Raw data and MATLAB code

Raw data and MATLAB analysis code used to generate all figures in this paper are deposited in the Dryad repository and are openly available via: http://dx.doi.org/10.5061/dryad.39h5m [55].

## Acknowledgements

The authors thank: Becky Ward, Orna Dahan, Justin Meyer, Matthew Bennett, and Frank Poelwijk for critically reading the manuscript; Anna Green and John Ingraham for help with preliminary experiments; Christine Degennaro, Sarah Boswell, Hana El-Samad, and Roy Kishony for helpful discussions; Shervin Javadi and Stratedigm Inc. for flow cytometry assistance.

## Abbreviations

GAL: galactose-utilization
YFP: yellow fluorescent protein
GEV: GAL4-DNA-Binding-domain.Estrogen-Receptor.VP16-activation-domain
MAL: maltose-utilization

## Supporting Information

**Text S1. All Supporting Figures and captions.**

Contains all the supporting figures and captions in one document.

**Table S1. Strains used in this study.**

This file contains three worksheets. Worksheet **(A)** lists the 43 natural isolates assayed by growth curves in Figure 1. Worksheet **(B)** lists the GAL1pr-YFP reporter strains constructed from a subset of 15 natural isolates. Worksheet **(C)** lists strains used in time-lapse microscopy and synthetic GAL induction experiments.

**Table S2. Phenotypic measurements of natural isolates.**

This file contains four worksheets. Worksheet **(A)** summarizes the metrics used in the paper and describes how they are defined and inter-related. Worksheet **(B)** contains values of the diauxic lag time and minimum mid-diauxic growth rate metrics from both replicates of the growth curve experiments in Figure 1. Worksheet **(C)** contains values of preparation time and other metrics measured on a subset of 15 natural isolates and used in Figures 3-4. Worksheet **(D)** contains growth rate measurements used to determine the galactose cost, as well as GAL expression data, plotted in Figures 7 and S14.

**Figure S1. Growth curves of all 43 strains assayed.**

Plots of log_2_ (OD_600_) versus time for 43 strains, after subtracting background (0.03) from the raw OD_600_ readings. Two replicates are shown in each panel. Time axes have been adjusted so that OD = 2^-6^ at time zero, to exclude an initial interval of 0-12 hours during which data can be noisy due to low OD (examples shown in Figure S2B). The strains are shown sorted from shortest to longest diauxic lag time from top left to bottom right. Plots with an asterisk “*” in top-right corner are strains shown in Figure 1B-C based on their galactose growth rate (Figure S3). Plots outlined in green represent the 15-strain subset used for GAL induction measurements (Table S1; Figure 3-4).

**Figure S2. Diauxic lag and minimum mid-diauxic growth rate metrics correlate across replicates.**

**(A)** Measured optical density (i.e. absorbance at 600nm) versus actual culture density, obtained by serial dilution of a yeast culture saturated under growth curve assay conditions. Dilution series were prepared in triplicate. OD600 was linear with culture density in this range, and displayed a background (y-intercept) value of ∼0.03. **(B)** Example growth curves (*top*) and growth rate plots (*bottom*) for 2 strains. Light gray lines show raw discrete derivatives computed from the growth curve data (Materials and Methods), blue lines show cubic spline fits to the discrete derivatives, and red lines show derivatives of the splines, or the smoothed 2^nd^ derivative of the growth curves. Both replicates are shown for each strain. Strain Bb32 (*left*) did not have a local growth rate minimum, and therefore its diauxic lag duration was defined to be zero and its minimum mid-diauxic growth rate was defined to be the time of the inflection point in growth rate (Materials and Methods). More often, strains displayed a phenotype like SLYG78 (*right*), a S288C derivative, which had a clear minimum rate during diauxic shift. **(C)** Scatter plots of the diauxic lag time (*left*) and minimum mid-diauxic growth rate (*middle*) across 2 replicate experiments, and between diauxic lag time and minimum rate (*right*). All 3 plots are strongly correlated, showing that our metrics were robust to measurement noise and that the continuous phenotypic variation in diauxic growth is not an artifact of the lag time metric used in Figure 1. **(D)** Diauxic lag time and minimum mid-diauxic growth rate were calculated from a sliding-window average on the discrete derivatives of growth curves (as opposed to the cubic spline fit used for Figure 1). This method yielded almost identical results, showing that the metrics were not sensitive to the method of calculation.

**Figure S3. Diauxic lag time is not correlated with growth rate in glucose-only or galactose-only medium.**

Scatterplots of diauxic lag time (*top*) and minimum mid-diauxic growth rate (*bottom*) versus steady-state growth rates in galactose alone (*left*) or in glucose alone (*right*). Steady-state growth rates were measured in a separate growth curve experiment (Materials and Methods). In general, the diauxic lag metrics correlated poorly with steady-state growth rates, suggesting that the phenotypic variation in diauxic growth cannot be solely explained by differences in glucose or galactose metabolism. Strains with growth rates between 0.5 and 0.6 doublings/hour in galactose are shown in Figure 1B-C (filled dots). This includes BC187 (blue) and YJM978 (red).

**Figure S4. Determination of absolute cell concentration by flow cytometric counting.**

**(A)** Absolute cell concentration as measured by flow cytometer versus actual relative culture density of a dilution series of a yeast culture, prepared in triplicate. Gray line shows predicted results extrapolated from the lowest density measurement, which agrees well with observed values. Absolute cell concentration was determined during the diauxic growth experiments in Figure 2B-C and 5A-B, and are plotted versus time for **(B)** BC187, **(C)** YJM978, and **(D)** a co-culture of BC187 and YJM978. Data for both replicates are shown—these are almost overlapping. The time axis is adjusted so that the culture is at 10^6^ cells/mL at time zero. This time was determined by interpolation on a linear fit to four consecutive datapoints. The same adjustment was applied to the time axis for all plots in Figures 2 and 5.

**Figure S5. Strain BC187 can consume galactose and glucose simultaneously.**

**(A)** Glucose and galactose concentrations versus time for (*left*) a culture of BC187, and (*right*) a culture of YJM978, from the same experiment as in Figure 2. Both replicates are plotted. Time zero corresponds to culture density of 10^6^ cells/mL. To determine whether either strain begins to consume galactose prior to glucose exhaustion, we computed **(B)** the sugar depletion rate by taking discrete derivatives (circles) of the sugar concentrations, or slopes between consecutive data points, for both replicate datasets. We then binned the time axis into 1-hour intervals and computed, via a one-tailed t-test, the probability of observing the discrete derivatives in each time interval given a null hypothesis that the mean discrete derivative in that interval is 0 or positive. Mean sugar depletion rate for each bin is shown as lines in (B) and the log_10_ p-value for the significance test is shown in **(C)**. Dotted black lines in (C) indicate where p=0.05. For BC187, there is a 2-hour interval over which there is statistically significant depletion of both glucose and galactose, at a significance threshold of 0.05. By contrast, the time intervals of significant glucose and galactose depletion for YJM978 do not overlap temporally. These conclusions are robust to the width or position of time bins.

**Figure S6. GAL1pr-YFP expression is highly variable across natural isolates in glucose + galactose but not in glucose alone.**

Steady-state GAL1pr-YFP expression histograms for 15 strains showing partial expression in 0.0625% glucose + 0.25% galactose (black), basal expression in 2% glucose (purple), and maximal expression in 2% galactose (orange). Additionally, the parent strains without a GAL1pr-YFP reporter cassette were assayed in 2% glucose (red). Partial expression varies widely across strains in glucose + galactose, yet YFP signal above autofluorescence is undetectable from most strains in glucose-only medium (compare purple and red histograms). A number of strains (YJM981, Y12-WashU, Y9-WashU, YJM975) display bimodal expression in glucose+galactose. Measurements were taken at steady-state (as in Figure 4D; see Materials and Methods); distributions are unsmoothed histograms of 20,000 or more cells.

**Figure S7. Co-culture method to determine timing of GAL induction relative to glucose depletion.**

**(A)** Example scatterplot of YFP versus mCherry signal by flow cytometry. Reference strain and query strain cells are distinguished by mCherry (red vs gray) and the YFP of each subpopulation is used to compute induction time. **(B)** Median GAL1pr-YFP expression of YJM978 with or without constitutive fluorophore in a co-culture of the two strains. Both strains contain the GAL reporter, which is unaffected by the constitutive fluorophore. **(C)** Start time of GAL induction t_low_ in BC187yb and YJM978ym cultured alone or in co-culture with each other, mean and range of 2 replicates. There is no significant difference in induction timing between separate and mixed cultures. **(D)** Median GAL1pr-YFP profiles for 15 strains from the co-culture experiment of Figure 3. Query and reference strain were mixed at three initial ratios. Density plot in background shows full YFP distributions of the query strain for the 1:4 query:reference condition (except for strain SLYG78, where 4:1 is shown). **(E)** Scatterplot of preparation time from different inoculating ratios. Preparation time was nearly identical across different inoculating ratios. The three ratios were used as replicates in Figure 3D-E. **(F)** Scatterplot of preparation time calculated as the time difference between query and reference strains at 1/32 or at 1/16 of maximal induction. The metric is robust to this difference (Spearman correlation = 0.97). **(G)** Definition of “ON fraction” as the fraction of cells with YFP signal higher than 1/32 of the maximal median YFP. **(H)** Possible ON fraction profiles. If a single population completely induces from basal to maximal (“Coherent induction”), the ON fraction will increase monotonically from 0 to 1. If the a culture splits into subpopulations with different induction times (“Early & late subpopulations”), the ON fraction will in two distinct phases, as in Venturelli et al. [23]. If a subset of cells never induce (“Non-inducing subpopulation”), the ON fraction will saturate below 1. **(I)** ON fraction versus time of the 15 strains in (D), from the 1:4 inoculation. Each strain is a different colored line, and strains BC187 and YJM978 are highlighted. Most profiles are consistent with “coherent induction”, although in some strains, a small subpopulation consisting of less than 10% of all cells may have pre-induction before sampling. In some strains, the ON fraction decreases after saturating (see also panel D) – this is likely due to an experimental artifact (Materials and methods).

**Figure S8. GAL1pr-YFP expression reaches steady-state after 8 hours of growth in galactose medium.**

GAL1pr-YFP expression distributions over time in repressing (0.25% glucose), inducing (0.25% galactose), and mixed-sugar (0.0625% glucose + 0.25% galactose) conditions for BC187yb (blue) and YJM978ym (red). Cultures were pre-grown in 2% raffinose to minimize the induction delay upon starting the experiment. Cells were diluted every two hours to maintain a density of less than 10^5^ cells/mL. After 12 hours, the dilution factor was increased and dilution / sampling interval increased to 12 hours, and the cultures were monitored up to 48 hours. In conditions where either strain induces, expression stops increasing after eight hours.

**Figure S9. Strains induce GAL1pr-YFP at quasi-steady-state levels during gradual glucose depletion.**

**(A)** Scatterplot of median GAL1pr-YFP expression of query strains three hours before reference strain mid-induction time (computed from data in Figure 3) versus the median GAL1pr-YFP expression of the same strains growing at steady-state in 0.0625% glucose + 0.25% galactose. **(B)** Steady-state GAL1pr-YFP distributions for strains BC187 and YJM978 (*Bottom*) in glucose + galactose conditions chosen from different moments of diauxic growth (*Top schematic*). BC187 induces at intermediate levels at steady-state in glucose + galactose mixtures, rather than at basal or maximal levels, as would be expected if the level of GAL expression responds in a switch-like manner to decreasing glucose.

**Figure S10. Pre-growth of YJM978 in a non-inducing galactose concentration accelerates GAL induction in subsequent medium shift.**

Median GAL1pr-YFP expression versus time for YJM978ym cells suddenly transferred from glucose to galactose (purple), or from glucose + galactose to galactose (black). This strain induces GAL genes significantly earlier (p=0.008 by 2-sample t-test) in response to sudden galactose induction when pre-grown in the presence of some galactose.

**Figure S11. Short-lag strains reach saturation faster, but BC187 exhausts glucose more slowly than YJM978.**

**(A)** Example calculation of saturation time, which is defined as the time for a strain to grow from the diauxic shift to saturation. **(B)** Scatterplot of saturation time versus diauxic lag time. The two metrics are strongly correlated, showing that strains that have a shorter diauxic lag also reach saturation sooner after diauxic shift. **(C)** Time to exhaustion of glucose or galactose in cultures of BC187yb (blue) or YJM978ym (red). YJM978 exhausts glucose significantly before BC187, even though BC187 exhausts galactose—and therefore both sugars—much faster than YJM978 under the assay conditions. Data are mean and range of n=2 replicates. P-value was calculated by 2 sample t-test.

**Figure S12. Absolute and relative fitness of BC187 and YJM978.**

**(A)** Log2 absolute cell concentration versus time of strains BC187yb (blue) and YJM978ym (red) in 0.0625% glucose (left) and 0.0625% glucose + 0.25% galactose (right). Cultures were sampled every two hours after they had reached steady-state GAL1pr-YFP expression. Cultures were periodically diluted so that raw cell densities (light color) did not exceed 2^20^ or 10^6^ cells/mL. Dilution-corrected data (dark color) were used to calculate growth rates. **(B)** Log2-ratio of BC187yb cell count to YJM978ym cell count in the same cultures as shown in (A). Relative fitnesses (i.e. growth rate differences) reported in Figure 5C are computed from line fits to these plots. **(C)** Steady-state growth rates of BC187yb (blue) and YJM978ym (red) in 0.0625% glucose + 0.25% galactose, 0.0625% glucose, or 0.15% galactose, as determined by linear fit to plots as in (A). Bar graphs are mean and s.e.m. of 3-6 replicates. P-values are computed by 2-sample t-test; “n.s.” indicates p > 0.05. **(D)** Steady-state GAL1pr-YFP expression distributions of BC187 (blue lines) and YJM978 (red lines) in the conditions from (C). Only one timepoint and replicate is shown; others had identical fluorescence distributions.

**Figure S13. Single-cell growth rate correlates negatively with GAL1pr-YFP expression**

**(A)** Example time-lapse microscopy images of BC187ym microcolonies (1-10 cells) at initial (top) and final (bottom) timepoints. Segmentation boundaries (red) were determined by analyzing mCherry images, and GAL1pr-YFP reporter concentration was determined as the final average YFP signal per pixel of each microcolony (Materials and methods). **(B)** Scatterplots of growth rate versus YFP concentration for microcolonies pre-induced in 0.125% glucose (n=165 microcolonies), 0.125% glucose + 0.25% galactose (n=196), or 0.25% galactose (n=223) prior to transferring to 0.125% glucose + 0.25% galactose for imaging. Growth rate displayed a significant negative correlation with GAL1pr-YFP concentration. **(C)** Distributions of growth rate (top) and YFP concentration (bottom) across microcolonies from the 3 pre-growth conditions. For clarity, plotted are smoothed probability densities estimated using a Gaussian kernel (Materials and methods). P-values are computed by a Kolmogorov-Smirnov test against the null hypothesis that growth rates of microcolonies from two pre-growth conditions have the same distribution.

**Figure S14. Galactose cost and GAL expression change in a correlated way between different media conditions.**

Scatterplot of galactose cost versus mean GAL1pr-YFP expression at steady-state in two different sets of glucose or glucose + galactose media. Black circles are the same data as in Figure 7B, whereas red circles are data obtained from 0.125% glucose and 0.125% glucose + 0.25% galactose, which induces GAL genes to a lesser extent. Gray arrows connect strains between the 2 conditions. Most arrows point toward the lower left, indicating that as galactose stays constant and glucose decreases (such as during diauxic growth), GAL expression increases at the same time that the growth cost due to the presence of galactose increases.

